# Susceptibility to Penicillin G and Ceftriaxone in Three Clinical *Treponema pallidum* Isolates is not Altered by Amino Acid Polymorphisms in the Tp0705 Penicillin Binding Protein

**DOI:** 10.1101/2025.10.14.682455

**Authors:** Lauren C. Tantalo, Kathyani D. Chamakuri, Alexander L. Greninger, Nicole A.P. Lieberman, Lorenzo Giacani

## Abstract

We demonstrated no differences in susceptibility to penicillin G and ceftriaxone in three modern *T. pallidum* isolates (UW244B, UW249B, and UW330B), each carrying a variant of the penicillin-binding protein (PBP) Tp0705. This suggests that these polymorphisms should not be a reason for concern when β-lactams are prescribed for syphilis treatment.

**SUMMARY:** We report that three modern *Treponem*a pallidum isolates carrying amino acid polymorphisms at positions 506, 625, and 708 of the penicillin-binding protein (PBP) Tp0705 are equally susceptible to penicillin G and ceftriaxone.

## INTRODUCTION

Syphilis, caused by the spirochete *Treponema pallidum* subsp. *pallidum* (*T. pallidum*), remains a global health concern, with a rising incidence worldwide (1, 2) and significant challenges in disease control. Once a syphilis diagnosis is established, the recommended treatment is penicillin G in different formulations and dosages, depending on disease stage and invasion of the central nervous system (3). In addition to penicillin G, other natural or synthetic β-lactam antibiotics have been shown to be effective against *T. pallidum* in pre-clinical (4) and clinical studies. In observational studies, for example, amoxicillin was highly effective in treating early syphilis (5, 6). Ceftriaxone, a potent third-generation β-lactam cephalosporin, is listed as a possible alternative to penicillin G to treat early syphilis, and data are available to support its efficacy for neurosyphilis (7-9). Promising results are also available on the efficacy for early syphilis treatment for cefixime, another third-generation cephalosporin (10, 11), although more studies are needed.

The reliance on β-lactams for syphilis treatment is justified by the lack of reliable evidence of resistance to this class of antibiotics after eight decades of use, consistent with their exceedingly low minimal inhibitory concentration (MIC) when tested *in vitro* against *T. pallidum* (4, 12). In contrast, there was rapid development and spread of resistance to macrolides when oral azithromycin was introduced for syphilis treatment (13). The *T. pallidum* genome does not contain plasmids or other mobile elements, which makes the acquisition and retention of environmental DNA coding for *bona fide* β-lactamases or other antibiotic resistance genes very unlikely. Nonetheless, we should not dismiss the possibility that amino acid polymorphisms on the pathogen penicillin-binding proteins (PBPs) could decrease the efficacy of β-lactams or even induce resistance, as hypothesized by Cha *et al*. (14) or by Mi *et al*. (15). These investigators, however, could not demonstrate their hypotheses due to lack of isolates with the proper genotypes to study. Furthermore, in 2025, two laboratory-derived mutant *T. pallidum* strains carrying the A1873G (inducing the amino acid change M625V) mutation in the PBP-encoding *tp0705* gene were reported to be partially resistant to ceftriaxone and penicillin G when exposed to low concentrations of these antibiotics (16), albeit with modest absolute effect sizes. Here, we tested the *in vitro* susceptibility to penicillin G and ceftriaxone of three *T. pallidum* clinical isolates (UW244B, UW249B, and UW330B) attained from patient blood samples, each carrying a distinct variant of Tp0705, due to polymorphisms at positions 1516, 1873, and 2122 of the gene, corresponding to amino acid position 506, 625, and 708 of the Tp0705 protein (Table 1) to see whether A1873G/M625V and other naturally occurring polymorphisms on the Tp0705 protein would alter susceptibility to β-lactams.

**Table 1.**
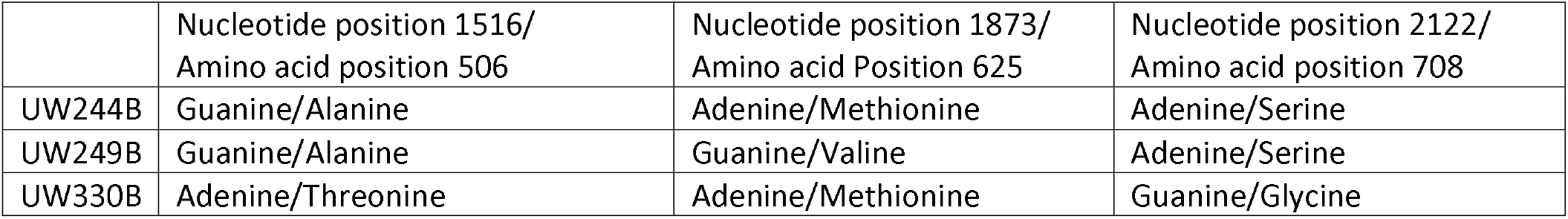
Polymorphisms in the *tp0705* gene/Tp0705 protein in the tested strains.

## METHODS

### *T. pallidum* strains and *in vitro* propagation

The UW244B, UW249B, and UW330B *T. pallidum* strains were isolated in 2004 (UW244B and UW249B) and 2005 (UW330B) by Dr. Christina Marra at the University of Washington via rabbit intratesticular injection with whole blood from patients with secondary (UW249B), early latent (UW330B), and late latent/unknown duration (UW244B) infection (17). For this study, all strains were grown *in vitro* as previously described (18) to perform antibiotic susceptibility testing. Genome sequencing data are available under BioProject/ID PRJEB20795/ERR2756217 (UW244B), PRJNA313497/SRR3268758 (UW249B), and PRJEB20795/ERR2756199 (UW330B) in INSDC databases. Polymorphisms were also confirmed by amplification/Sanger sequencing of the *tp0705* gene (see Supplementary Methods). The UW244B, UW249B, and UW330B isolates are related to SS14-clade strains (n=1582 sequenced genomes) from each continent by a median of 10 or fewer core genome SNPs and are therefore representative of circulating strains (17).

### Penicillin G and ceftriaxone susceptibility testing

Susceptibility testing of the three selected strains was performed as previously described (4) by incubating *in vitro* for seven days *T. pallidum* with increasing concentrations of penicillin G or ceftriaxone above and below the MIC previously determined for these antibiotics (4, 19). Concentrations ranged from 0.125-12 ng/ml (penicillin G), or 0.625-25 ng/ml (ceftriaxone), followed by determination of treponemal growth by qPCR in test and control (i.e., no antibiotic and antibiotic diluent only) replicate culture wells. In both assays penicillin G at 60 ng/ml was used as a treponemicidal control. Penicillin G (Cat. #J63032) and ceftriaxone (Cat. #C5793) were purchased from Thermo Fisher (Waltham, MA) and Millipore Sigma (Burlington, MA), respectively. The susceptibility assay is described in more detail in Supplementary Methods. Two-way ANOVA test was used to calculate statistical significance (\**p*⍰= 0.01-0.05) among medians within a concentration group.

## RESULTS

Incubation of the three strains with penicillin G resulted in complete killing of *T. pallidum* when exposed to a concentration of 1.00 ng/ml or higher of antibiotic (Fig.1A). Below 1.00 ng/ml, some treponemal growth occurred, although not as efficiently as when the antibiotics were absent from the culture media, consistent with previous reports that identified the secondary MIC for penicillin G (i.e., the lowest antibiotic dilution at which growth occurred but at a significantly lower rate compared to the no-antibiotic control wells) to be 0.5 ng/ml (19). In one instance (Fig.1A), we found a significant difference in growth between the UW249B and UW330B strains exposed to 0.25 ng/ml of penicillin G. This difference, however, was not confirmed at any other non-treponemicidal concentration tested. Similarly, exposure to ceftriaxone concentrations equal to or above 2.50 ng/ml was fully bactericidal *in vitro* (Fig.1B), which was also consistent with previous studies (4), while a single significant difference in growth between the UW244B and UW249B strains could be seen upon exposure to 0.625 ng/ml of antibiotic (Fig.1B), but not confirmed at any other concentration which was not fully bactericidal.

**Figure 1.**
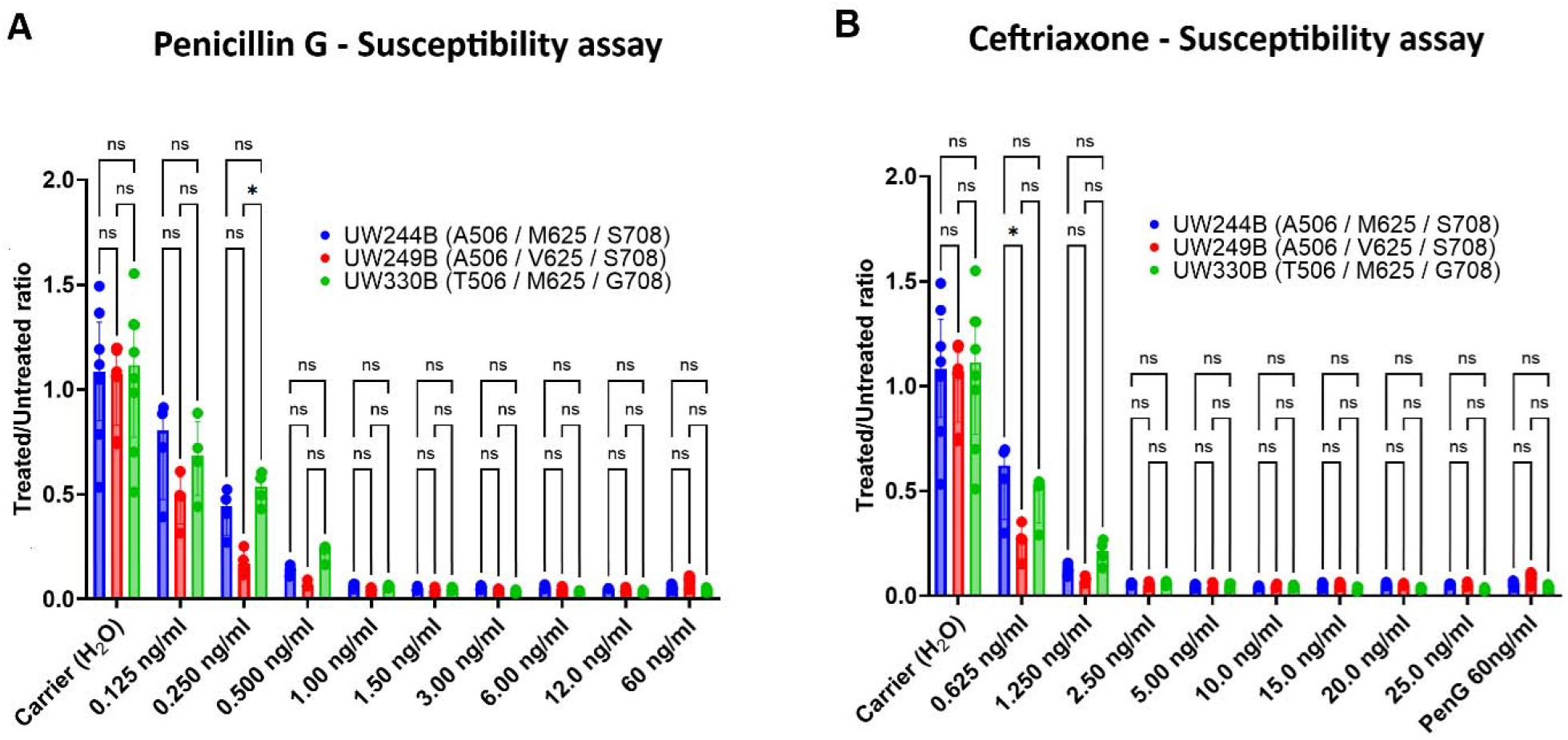
Susceptibility assay for penicillin G and ceftriaxone using the UW244B, UW249B, and UW330B isolates. Strain legend reports the amino acid polymorphism at positions 506 (Alanine or Threonine), 625 (Methionine or Valine), and 708 (Serine or Glycine), respectively, induced by nucleotide differences at positions 1516, 1873, and 2122 in the *tp0705* gene. All graphs are based on the number of *tp0574* gene copies detected in culture at Day 7 post-inoculation by qPCR. *tp0574* copies were normalized to the copies present in the corresponding controls (i.e., treponemes grown without antibiotics). Bars are the median ratio value +/-interquartile range (IQR), with dots representing individual ratio values. In all experiments, penicillin G at a concentration of 60 ng/ml was used as positive control for treponemicidal activity. Eight biological replicates were examined using qPCR for controls and four for test concentrations of penicillin or ceftriaxone. **(A)** Strain susceptibility to concentrations of penicillin ranging from 0.125 to 12 ng/ml. **(B)** Strain susceptibility to ceftriaxone concentrations ranging from 1.25 to 25 ng/ml. The two-way ANOVA test was used to calculate statistical significance (\**p*⍰= 0.01-0.05) among medians within a concentration group.

## DISCUSSION

Penicillin-binding proteins involved in cell wall synthesis are the targets of β-lactam antibiotics, and mutations/polymorphisms in PBPs can reduce susceptibility to penicillin or induce resistance by decreasing the affinity for the drug of the transpeptidase domain of these enzymes (20). Although *T. pallidum* has been treated with penicillin for decades, and penicillin resistance has never been experimentally demonstrated, rare reports of treatment failures following administration of β-lactams exist (15, 16, 21, 22). Given the pivotal role of β-lactams in syphilis treatment, evaluating whether allelic variants of the five known *T. pallidum* PBPs (the Tp0500, Tp0547, Tp0574, Tp0705, and Tp0760 proteins) confer differential susceptibility to β-lactams is a worthy task, now possible via a novel cultivation system (18).

The isolates analyzed here did not show significant differences in the penicillin G susceptibility assay (Fig.1A), although UW249B grew slightly less when exposed to 0.250 ng/ml of penicillin G. This difference, however, was not seen at immediately higher (0.5 ng/ml) or lower (0.125 ng/ml) concentrations, suggesting that the antibiotic concentration range at which this phenomenon is detectable might be very limited.

When a patient receives a single intramuscular (IM) injection of 1.2 M units of benzathine penicillin G (half the total dose a patient with early syphilis would receive) and the blood plasma concentration is measured daily over time, the recorded values range from 119 ng/ml (24 hours post-injection, corresponding to ∼238 times the penicillin G MIC for *T. pallidum*) to 5.0 ng/ml (at 29 days post-injection, corresponding to ∼10 times the MIC) (23). All concentrations in the 119-5.0 ng/ml range are highly treponemicidal for *T. pallidum in vitro*, as shown independently by us and other investigators (4, 19). Although no data on the concentration of penicillin G in tissue could be found after IM injection, even if the different Tp0705 alleles studied here could modulate susceptibility to penicillin G, this effect would manifest at antibiotic concentrations that are too low to be clinically relevant, given that one single 2.5M unit dose of benzathine penicillin G administered IM is sufficient to treat uncomplicated syphilis (24).

A similar conclusion can be drawn for ceftriaxone, for which a plasma concentration of 80.5 μg/ml (∼32,200 times the reported ceftriaxone MIC for *T. pallidum*) (4) and 29.7 μg/ml (∼11,880 times the MIC) are achieved by a single IM dose of 1 g (25) at 1 hour and 24 hours post-injection, respectively. Far higher plasma concentrations were reached by intravenous (IV) injection of ceftriaxone in the same experiment (25). In our study, slight differences in strain growth upon ceftriaxone exposure could be seen only at the lowest concentration tested (0.625 ng/ml; Fig.1B), although such differences were not confirmed at higher concentrations. Considering that ceftriaxone therapy for syphilis requires a 10-day daily administration regimen, either IM or IV, the difference seen here (Fig.1B) also seems negligible from a disease management standpoint.

Interestingly, the polymorphism associated with decreased susceptibility of the laboratory-derived strains used in (16), A1873G/M625V, did not seem to confer a similar advantage to strain UW249B, also carrying the same polymorphism, in our study. This, in our opinion, reiterates the importance of using clinical isolates in assessing drug susceptibility whenever possible. In the past, we also tested the susceptibility to ceftriaxone of the historical SS14 strain that, like the widely used Nichols strain, carries the A506, M625, and G708 polymorphisms in Tp0705 (4). These earlier data showed that for this strain, like the ones studied here, a ceftriaxone concentration equal or above 2.5 ng/ml is treponemicidal.

Furthermore, we recently performed an azithromycin susceptibility experiment using the UW330B and SS14 isolates, both known to be macrolide resistant via different mutations (A2058G or A2059G) in the 23S rRNA gene (4). These experiments showed that when exposed to 64 times the azithromycin MIC for *T. pallidum*, these isolates grew as if no antibiotic was present in the media, further supporting that, while UW330B and most of the modern syphilis strains circulating worldwide are indeed fully resistant to azithromycin (13), the same cannot be said for resistance to β-lactams in the isolates tested in this study based on our experimental results.

## ACKNOWLEDGEMENTS

This study was supported by NIH Grant R56AI175016 (to L.G.). The funders of the study had no role in the study design, data collection, data analysis, data interpretation, manuscript writing, or in the decision to submit it for publication. We are thankful to Dr. Christina Marra for providing the *T. pallidum* UW strains tested in this study, and to Dr. Jodie Dionne (UAB-Birmingham, AB, USA) for her helpful comments on the manuscript ahead of submission.

## REFERENCES

1. WHO. Global progress report on HIV, viral hepatitis and sexually transmitted infections, 2021 - Accountability for the global health sector strategies 2016–2021: actions for impact. https://wwwwhoint/publications/i/item/9789240027077. 2021.

2. WHO. World Health Organization. Syphilis. 2024. https://wwwwhoint/news-room/fact-sheets/detail/syphilis. 2024.

3. CDC. STD Treatment Guidelines - Syphilis. Morbidity and Mortality Weekly Report. 2021.

4. Tantalo LC, Lieberman NAP, Pérez-Mañá C, Suñer C, Vall Mayans M, Ubals M, et al. Antimicrobial susceptibility of Treponema pallidum subspecies pallidum: an in-vitro study. Lancet Microbe. 2023;4(12):e994–e1004.

5. Nishijima T, Kawana K, Fukasawa I, Ishikawa N, Taylor MM, Mikamo H, et al. Effectiveness and Tolerability of Oral Amoxicillin in Pregnant Women with Active Syphilis, Japan, 2010-2018. Emerg Infect Dis. 2020;26(6):1192–200.

6. Tanizaki R, Nishijima T, Aoki T, Teruya K, Kikuchi Y, Oka S, et al. High-dose oral amoxicillin plus probenecid is highly effective for syphilis in patients with HIV infection. Clin Infect Dis. 2015;61(2):177–83.

7. Shann S, Wilson J. Treatment of neurosyphilis with ceftriaxone. Sex Transm Infect. 2003;79(5):415–6.

8. Cao Y, Su X, Wang Q, Xue H, Zhu X, Zhang C, et al. A Multicenter Study Evaluating Ceftriaxone and Benzathine Penicillin G as Treatment Agents for Early Syphilis in Jiangsu, China. Clin Infect Dis. 2017;65(10):1683–8.

9. Marra CM, Boutin P, McArthur JC, Hurwitz S, Simpson PA, Haslett JA, et al. A pilot study evaluating ceftriaxone and penicillin G as treatment agents for neurosyphilis in human immunodeficiency virus-infected individuals. Clin Infect Dis. 2000;30(3):540–4.

10. Stafylis C, Keith K, Mehta S, Tellalian D, Burian P, Millner C, et al. Clinical Efficacy of Cefixime for the Treatment of Early Syphilis. Clin Infect Dis. 2021;73(5):907–10.

11. Klementová T, Zákoucká H, Bížová B, Unemo M, Rob F. Cefixime versus benzathine penicillin G for the treatment of early syphilis-a randomized, controlled open label trial. J Antimicrob Chemother. 2025.

12. Hayes KA, Dressler JM, Norris SJ, Edmondson DG, Jutras BL. A large screen identifies beta-lactam antibiotics which can be repurposed to target the syphilis agent. NPJ Antimicrob Resist. 2023;1(1):4.

13. Lieberman NAP, Reid TB, Cannon CA, Nunley BE, Berzkalns A, Cohen SE, et al. Near-Universal Resistance to Macrolides of Treponema pallidum in North America. N Engl J Med. 2024;390(22):2127–8.

14. Cha JY, Ishiwata A, Mobashery S. A novel beta-lactamase activity from a penicillin-binding protein of Treponema pallidum and why syphilis is still treatable with penicillin. J Biol Chem. 2004;279(15):14917–21.

15. Mi HF, Shen X, Chen XQ, Zhang XL, Ke WJ, Xiao Y. Association between treatment failure in patients with early syphilis and penicillin resistance-related gene mutations of Treponema pallidum: Protocol for a multicentre nested case-control study. Front Med (Lausanne). 2023;10:1131921.

16. Pospíšilová P, Bosák J, Hrala M, Krbková L, Vrbová E, Šmajs D. Resistance to ceftriaxone and penicillin G among contemporary syphilis strains confirmed by natural in vitro mutagenesis. Commun Med (Lond). 2025;5(1):224.

17. Beale MA, Marks M, Sahi SK, Tantalo LC, Nori AV, French P, et al. Genomic epidemiology of syphilis reveals independent emergence of macrolide resistance across multiple circulating lineages. Nat Commun. 2019;10(1):3255.

18. Edmondson DG, Hu B, Norris SJ. Long-Term In Vitro Culture of the Syphilis Spirochete Treponema pallidum subsp. pallidum. mBio. 2018;9(3).

19. Norris SJ, Edmondson DG. In vitro culture system to determine MICs and MBCs of antimicrobial agents against Treponema pallidum subsp. pallidum (Nichols strain). Antimicrob Agents Chemother. 1988;32(1):68–74.

20. Nagai K, Davies TA, Jacobs MR, Appelbaum PC. Effects of amino acid alterations in penicillin-binding proteins (PBPs) 1a, 2b, and 2x on PBP affinities of penicillin, ampicillin, amoxicillin, cefditoren, cefuroxime, cefprozil, and cefaclor in 18 clinical isolates of penicillin-susceptible,-intermediate, and -resistant pneumococci. Antimicrob Agents Chemother. 2002;46(5):1273–80.

21. Kenyon C, Vanbaelen T. Penicillin treatment failure probably explains repeat episode of neurosyphilis. The Lancet Infectious Diseases. 2025;25(4):e194.

22. Tramont EC. Persistence of Treponema pallidum following penicillin G therapy. Report of two cases. Jama. 1976;236(19):2206–7.

23. Broderick MP, Hansen CJ, Russell KL, Kaplan EL, Blumer JL, Faix DJ. Serum penicillin G levels are lower than expected in adults within two weeks of administration of 1.2 million units. PLoS One. 2011;6(10):e25308.

24. Hook EW, 3rd, Dionne JA, Workowski K, McNeil CJ, Taylor SN, Batteiger TA, et al. One Dose versus Three Doses of Benzathine Penicillin G in Early Syphilis. N Engl J Med. 2025;393(9):869–78.

25. Goonetilleke AK, Dev D, Aziz I, Hughes C, Smith MJ, Basran GS. A comparative analysis of pharmacokinetics of ceftriaxone in serum and pleural fluid in humans: a study of once daily administration by intramuscular and intravenous routes. J Antimicrob Chemother. 1996;38(6):969–76.

