## supplementary methods for "Susceptibility to Penicillin G and Ceftriaxone in Three Clinical *Treponema pallidum* Isolates is not Altered by Amino Acid Polymorphisms in the Tp0705 Penicillin Binding Protein": SUPPLEMENTARY METHODS.docx

#### T. pallidum strains

Live treponemes were retrieved from frozen stocks obtained through intratesticular strain propagation in New Zealand White rabbits (*Oryctolagus cuniculus*). Although no animals were used for the study described here, care for the animals used to propagate the study strains was provided according to the Guide for the Care and Use of Laboratory Animals. Animal procedures for those experiments were approved and covered by the University of Washington IACUC protocol # 4243-01 (PI: Lorenzo Giacani).

#### Cell culture and T. pallidum inoculation for susceptibility assay

For the susceptibility assay, we used columns of 96-well plates (Corning Inc., 8x12 format) to test different drug concentrations for each selected antimicrobial. Each drug concentration was considered as a separate experimental group and tested in four replicate wells within a column. Control groups where treponemes were grown in the absence of antibiotics, and treponeme cultures where the antibiotic was replaced by the antibiotic carrier (water). Controls were tested in eight replicate wells within a column.

Each antibiotic solution (1.5 µl) was added from a 100X concentrated stock to achieve the final concentration to be tested without significantly altering the final volume of the culture. After the addition of the treponemes and agents/solvents, the culture wells were incubated at 34°C in the tri-gas incubator until harvest. Treponemes from experimental and control wells with the selected antibiotic concentration range were harvested to perform DNA quantification after a week-long incubation.

The day before treponemal inoculation, wells (96-well plates) were seeded with 3x10^3^ rabbit Sf1Ep cells in 150 μL of MEM culture media. The plates were then incubated overnight in a 5% CO_2_ atmosphere within a HeraCell 150 incubator (Thermo Fisher Scientific, Waltham, MA) to allow Sf1Ep cell adhesion to the well surface. On the same day, TpCM2 treponemal media was prepared as previously reported ^1^ and equilibrated overnight in a HeraCell 150i tri-gas incubator (Thermo Fisher Scientific) at 34°C in a microaerophilic environment (1.5% O_2_, 3.5% CO_2_ and 95% N_2_). The following day, MEM was removed from the Sf1Ep-containing wells, and cells were rinsed with 150 µL equilibrated TpCM2 media. Subsequently, 150 µL of equilibrated TpCM2 media were added to each well, and the plates were placed in the tri-gas incubator for at least three hours. *T. pallidum* cells grown on Sf1Ep cultures inoculated the previous week were separated through trypsinization to allow the release and quantification of spirochetes and to prepare the inoculum for the 96-well test plates. Treponemes were counted using dark field microscopy on a Leica DM2500 LED microscope (Leica, Wetzlar, Germany) and diluted in TpCM2 to 3.3x10^5^ *T. pallidum* cells/ml.

#### Harvest and DNA extraction/quantification

The exhausted TpCM2 was removed from wells and discarded. A total of 200 µL of genomic lysis buffer (Zymo Research) was added to each well, and wells were incubated for 30 minutes at room temperature to allow for cell lysis as per the protocol provided by the kit manufacturer. Processed plates were sealed and stored at -20°C until DNA extraction. In earlier studies ^2^, we determined that most (~85%) of *T. pallidum* cells *in vitro* adhere to the rabbit epithelial cell monolayer, as also reported by Edmondson *et al.* for other cultivated strains ^3^. This evidence allowed us to discard the culture media without concern that the experimental results would be significantly affected.

##### DNA extraction and quantification

Treponemes were pelleted from each aliquot at 20,000 x g for 10 minutes and, after supernatant removal, pellets were resuspended in 200 µL of genomic lysis buffer provided with the Quick-DNA 96 kit (Zymo Research) and stored at -20⁰C until DNA extraction. To extract the DNA, the 96-well plates were thawed at 37°C and spun briefly to remove condensation drops on the plate sealers. DNA was extracted using the Quick-DNA 96 kit (Zymo Research) according to the manufacturer’s instructions. DNA was eluted in 100 µL of molecular water and stored at -20⁰C until analysis. DNA obtained from each sample was quantified by qPCR targeting the *T. pallidum*-specific *tp0574* gene as previously described ^4^. The Powerup SYBR Green Master Mix (Thermo Fisher Scientific) was used for amplification. Amplifications were run on a QuantStudio3 or QuantStudio5 thermal cycler (Thermo Fisher Scientific), and results were analysed using the instrument software. Data were imported into Prism 8 (GraphPad Software, San Diego, CA) and further analysed to assess the statistical significance of the values from test and no-antibiotic control groups using one-way ANOVA with the Dunnett test for correction of multiple comparisons with significance set at *p*<0.05 in both cases.

### Amplification and sequencing of the *tp0705* gene

The *tp0705* locus was amplified by PCR using DNA extracted from each strain used in theis study. The total volume of PCR reactions was 50 μL, with the following composition: 3 μL of template DNA, 1.5 μL of dNTP mix, 2.5 μL Mg2+ plus buffer and 0.05 μL Taq polymerase, 3 μL of each primer (0.025 M). Amplification of all tested loci was performed under the following cycling conditions: 94°C (1 min); 98°C (10 s), 68°C (15 s) touch down (−1.0°C per cycle), and 68°C (1 min, 45 s) for 8 cycles; 98°C (10 s), 61°C (15 s), and 68°C (1 min, 45 s) for 35 cycles, with final extension at 68°C (7 min). PCR products were purified and sequenced by Sanger sequencing by Azenta. Sequence analyses were performed using the Bioedit software and merged in a gene alignment for each locus to check each sequence and obtain its corresponding allele. Amplification and sequencing primers are reported below. Amplicon lenght was of 1181 bp.

**Primers used for amplification and sequencing of *tp0705***

| Amplification/Sequencing primers (5-3) | Coordinates^1^ | Seqeuncing primers (5-3) | Coordinates^1^ |
| --- | --- | --- | --- |
| GGTCTATATGCAGCCCTTCTTC | 772663-772684 | TGCGGCTTATCCTGATGAATAG | 772917-772938 |
| GCTTGAGAACGATACCGGATAC | 773822-773843 | TATTCTGCGGCGTTGGATAG | 773700-773719 |

^1^Based on the Nichols genome (CP004010.2)

1.	Phan A, Romeis E, Tantalo L, Giacani L. In Vitro Transformation and Selection of Treponema pallidum subsp. pallidum. *Curr Protoc* 2022; **2**(8): e507.

2.	Haynes AM, Giacani L, Mayans MV, et al. Efficacy of linezolid on Treponema pallidum, the syphilis agent: A preclinical study. *EBioMedicine* 2021; **65**: 103281.

3.	Edmondson DG, Hu B, Norris SJ. Long-Term In Vitro Culture of the Syphilis Spirochete Treponema pallidum subsp. pallidum. *mBio* 2018; **9**(3).

4.	Giacani L, Molini B, Godornes C, et al. Quantitative analysis of tpr gene expression in Treponema pallidum isolates: Differences among isolates and correlation with T-cell responsiveness in experimental syphilis. *Infect Immun* 2007; **75**(1): 104-12.
